# A versatile survey method for repeatedly monitoring individual road-kill

**DOI:** 10.1101/2025.05.06.652546

**Authors:** Jon J Sullivan, Giorgia Vattiato, Chris N Niebuhr

## Abstract

Road-kill is increasingly recognised as an important source of mortality for wildlife, especially in densely populated urban and rural landscapes. Monitoring road-kill on fine spatial and temporal scales is necessary to better understand how the design of road networks affects road-kill. We report on a practical method for consistently and repeatedly surveying road-kill. This includes geotagging individual carcasses, reporting where on or by the road each carcass is found, categorising the age and state of each carcass, and noting whether each carcass is new or previously reported. All this can practically be done with a smartphone while on foot, biking, or as a passenger in a motor vehicle. Repeatedly mapping all carcasses, whether or not they have previously been reported, allows for estimates of carcass persistence and detection probability for both taxa and road conditions. We demonstrate the effectiveness of the method by reporting on its frequent use along one 16.2 km route of urban and rural roads in New Zealand over 12 years. Over this period, on 1,652 surveys, 4,034 new road-kill carcasses were observed, 62.4% of which were birds, 31.1% were mammals, and 3% were butterflies. Using carcass age and persistence, we estimate that the road-kill rate along this route has been at least 3,940-6,544 road-kill/100 km/year. There was considerable variation among taxa in carcass persistence and carcass position on and by the road, both of which will introduce biases into road-kill estimates if not accounted for. To better understand road-kill and estimate road-kill rates, we encourage road-kill studies to geotag all individual carcasses and track their persistence.

## Introduction

Vehicle collisions result in several hundred million bird mortalities worldwide each year, including estimates of 14 million in Canada, 27 million in England, and 80–340 million in USA (as cited in Husby, 2017). In New Zealand, much of the reported road-kill is of smaller, introduced mammals, e.g. common brushtail possums (*Trichosurus vulpecula*), hedgehogs (*Erinaceus europaeus*), European rabbits (*Oryctolagus cuniculus*) and hares (*Lepus europaeus*), as opposed to ungulates, resulting in little public concern regarding risk to motorists. Limited information exists on the impacts of vehicle-caused mortality on avian species or populations in New Zealand, although some reports exist that provide evidence of birds being killed by vehicles and highlight risks to populations and individuals (e.g. Beauchamp, 2009; Brockie, Sadleir & Linklater, 2009; Sadleir & Linklater, 2016; Flux, Tryjanowski & Zduniak, 2022). We have only identified eight studies within the peer-reviewed scientific literature that are primarily dedicated to road-kill in New Zealand. For these studies (which are discussed in detail in the following section), reporting, survey methods, sampling designs, and analyses differed, making simple comparisons difficult.

Direct comparisons with the international literature may be even more difficult, due to differences in New Zealand avifauna. For example, some of New Zealand’s endemic bird species have evolved flightlessness in the historical absence of mammalian predators, likely increasing their vulnerability to road mortality. Furthermore, these studies were typically not conducted within major habitats of highly threatened species, and thus present data highly dominated by more common native and introduced species (e.g. Sadleir and Linklater 2016 conducted surveys in peri-urban and farmland habitats). Currently 80 bird species in New Zealand are classified ‘Threatened’ and 98 as “At Risk” by the NZ Threat Classification system used by the Department of Conservation (Robertson et al., 2021).

Mitigation efforts are designed to reduce mortality/injury due to road-kill. Over 40 types of road mitigation, designed to reduce road-kill of wildlife, exist globally, and these are discussed in detail in various reviews and reports (e.g. Huijser et al., 2008; van Der Ree, Smith & Grilo, 2015; Rytwinski et al., 2016). There is no ‘one size fits all’ mitigation practice, and each site and/or species scenario must be individually considered. Regardless of the type of mitigation used, proper evaluation of the effectiveness of any effort should include road-kill data collected prior to implementing the to allow comparisons to data collected after the mitigation attempts have taken place. In order to collect such data, an appropriate sampling method must be available that accurately estimates road-kill numbers while minimising and quantifying any biases.

Multiple methods have been used to monitor road-kill of a given species or in a certain area (van Der Ree, Smith & Grilo, 2015). Some monitoring is passive (often opportunistic and without an experimental design), such as police or road agency reports and road-kill citizen science efforts. Other monitoring is active (implementing a predetermined, experimental design), such as government mandated surveys (often poorly designed and under-resourced) and research-based surveys (question-driven and usually more rigorous) (van Der Ree, Smith & Grilo, 2015). Regardless of the type of survey method used, there is often bias associated with the data collected, such as overrepresentation of certain species or underestimations of true road-kill numbers occurring in the landscape, which makes comparing data difficult (Guinard, Julliard & Barbraud, 2012). Unbiased estimates of road-kill abundance are necessary to accurately assess the impacts on a population, community, or species, as well as measure the effectiveness of any mitigation measures designed to reduce these impacts. One review reported that out of 645 publications discussing research issues related to wildlife vehicle collisions, only 7% dealt with the approach of data collection (Pagany, 2020). (Guinard, Prodon & Barbraud, 2015) suggested that to reduce bias in road-kill surveys, three parameters need to be quantified:

– Persistence (the probability that the carcass was not removed from the road between counts; or the inverse of carcass disappearance)
– Entry (the probability that a new carcass appears on the road between counts)
– Detectability (the probability of detecting the carcass)

Very few peer-reviewed studies have reported on road-kill of birds in New Zealand, with even fewer having conducted designed sampling. Of the eight New Zealand studies with road-kill as the primary focus (Burger & Gochfeld, 1992; Brockie, 2007; Washington et al., 2008; Beauchamp, 2009; Brockie, Sadleir & Linklater, 2009; Freeman, 2010; Sadleir & Linklater, 2016; Flux, 2019), only five focussed on multiple species. Seven studies conducted driving counts, with one conducting cycling counts.

Here we present a versatile, consistent method to ascertain rates of road-kill that has been applied over the past two decades, mostly while travelling by bike, in part of New Zealand. We demonstrate that it is practical to track individual road-kill carcasses on daily work-commutes over extended periods of time with relatively little effort, in a way that allows road-kill detection and persistence biases to be estimated.

## Materials & Methods

Our method allows for the convenient collection of individually geotagged road-kill while moving on foot, by bicycle, or as a passenger in a motor vehicle. It is designed to quickly and accurately map out all road-kill visible along survey routes. Dr. Jon Sullivan, Lincoln University, has used this method, or precursors of it, to collect almost daily road-kill data beginning in 2003, mainly around Ōtautahu-Christchurch, NZ (see case study below). At a minimum, the method requires a device recording a GPS track and a means of recording timestamped written or audio notes. All can be provided by a smart phone.

The survey is marked with a start and stop time, and then each road-kill along the route is marked with a timestamped note, either written in shorthand or spoken. This proved to be much faster data entry than more complex interfaces with drop-down options or buttons to browse through. There are many apps that can create timestamped notes. For audio notes, a simple voice recorder can be sufficient, so long as it accurately records the start time of each recording (e.g. Voice Memos). We use FileMaker Go for its flexibility in recording timestamped (and geotagged) text, audio, and handwritten notes. The TRYVL app for mobile devices is another option that geotags voice memos.

The timestamps of all road-kill observations can later be geotagged with the GPS track. We have found that geotagging against a continually recording GPS track provides substantially more accurate coordinates that relying on a smart phone’s point geotags. Points by a smart phone while moving along a road can lag by many tens of metres. The continuous GPS track can be made by a smart phone app (we use the Cyclemeter app) or using a standalone GPS unit (we use a Garmin GPSMAP 64).

Road-kill observations can be either written in shorthand or spoken using an analogous syntax. Each uses the following structure, where asterisked elements are compulsory and non-asterisked elements are optional:

[species name]* [identification uncertainty] [position on/by road]* [road-kill age]* [road-kill condition]* ([alternative identifications] [new or usual carcass] [other notes, e.g. sex, life stage])

For example, we could type “hh lr old sq (usual)” or speak “hedgehog left road old squashed usual”. Both indicate that at this location there was a road-kill carcass of a hedgehog (*Erinaceus europaeus*) on the left side of the road that was old (definitely >24 hours old) and squashed on the road. “Usual” indicates that it was a carcass that had been already recorded on a previous survey and was still present. We expand on the rationale and categories for each of these elements below.

### Road-kill taxon

Each road-kill observation begins with the taxon name. Typically this is a species name although it can be a higher taxon when the identification is difficult (e.g. raptor, bird, mammal). A user-maintained list matching short-hand names to scientific names is required (e.g. hh, *Erinaceus europaeus*).

When the observer is not 100% certain of a species identification, it is always followed by a question mark (e.g. “hh?” in shorthand, “hedgehog question mark” in spoken audio). In rare cases where it is not certain whether an object on the road is a road-kill, a double question mark is used (e.g. “hh??” in shorthand, “hedgehog question mark question mark” in spoken audio). This most often happens when the road-kill is well-worn and observations are made from a fast-moving car. This is important so that the overall road-kill density per kilometre is not influenced by the observer’s decisions around whether objects that look like well-worn carcasses are or aren’t dead animals.

### Road-kill age

Road-kill age is provided as “fresh” (definitely <24 hours old), “unsure” (possibly <24 hours old), and “old” (definitely >24 hours old). This helps with calculations of road-kills per day along regularly travelled routes. With some experience, especially along regularly surveyed routes, categorising road-kill to this coarseness of age is usually quite easy.

### Road-kill condition

The condition of a carcass on the road affects its detection probability and its persistence on the road (e.g. availability to scavengers). This is recorded as “intact”, “exposed” (some internal tissues visible but carcass still fully formed), “squashed”, “decomposed” (carcass visibly flattened down due to decomposition), “fragment” (only part of the tissue of the carcass remains), and “skeleton” (only bones are visible). In shorthand, these are abbreviated as “int”, “exp”, “sq”, “decomp”, “frag”, and “skel” respectively. Additional condition categories “fur”, “feathers”, and “blood” are optionally used although in our surveying only carcasses with tissue present have been consistently recorded.

### Road-kill position

Like condition, the position of the carcass on or by the road affects its detection probability and persistence. Carcasses on the road in the path of traffic are more apparent but less persistent. For simplicity, seven categories are used, “mid road” (within 30 cm of the road centre line), “left road” and “right road” (on the road in the path of vehicles, relative to the direction of the observer), “left verge” and “right verge” (either beyond the roadside boundary line but still on the road, or within 30 cm of the road edge away from the path of car tyres), and “left roadside” and “right roadside” (synonymous in our data collection with “left grass” and “right grass”).

These categories are abbreviated in shorthand as “mr”, “lr”, “rr”, “lv”, “rv”, “lg”, “rg” respectively. “Left road” and “right road” can be optionally replaced with compass directions for added clarity (“east road”, “west road”, “north road”, “south road”). This can be useful when on foot in complex intersections where “right” and “left” may be misinterpreted.

### Road-kill persistence

Understanding road-kill persistence and detection probability are both assisted by repeat observations of the same carcasses. In our method, all carcasses are recorded on every survey, regardless of whether or not they have been recorded previously. All previously recorded carcasses are noted with the special word “usual” (with care being taken not to use “usual” in other notes). When it is uncertain whether or not a carcass is usual, “possibly usual” and “probably usual” are used. For frequently repeated surveys, this knowledge of which carcasses are new and which are usual can be used to provide estimates of road-kill persistence for different taxa, in different conditions, on different positions on and by the road. That in turn can be used to better interpret the road-kill rates from data collected on less frequently travelled routes.

### Other notes

Other notes about a carcass are optionally added after the essentials. In shorthand, these notes are in parentheses. In spoken notes, they are just said after the essentials. When apparent, sex and life stage (“adult”, “juvenile”, “baby”) are noted. The word “photo” is exclusively used when a photo of a carcass is taken.

### Multiple carcasses at one location

When there are multiple carcasses at one location, each individual animal is always entered separately. For shorthand notes, this is done by putting each carcass on a separate line of one geotagged note. In spoken audio, each observation is separated by “and also with”. For example,

pd mr fresh int (female)

pd lr fresh sq (baby)

pd rr fresh sq (baby)

“paradise shelduck mid road fresh intact female and also paradise shelduck left road fresh squashed baby and also paradise shelduck right road fresh squashed baby”

This typed shorthand, or equivalent spoken note, indicates that the carcass of an adult female paradise shelduck was in the middle of the road and two paradise shelduck ducklings were squashed, one on the left side of the road and one on the right side.

### Processing audio notes

When passenger in a car, or walking, it is practical to type in road-kill as shorthand. Care needs to be taken with smart phones that their automatic spellcheckers do not override what is typed. Shorthand codes can be added to a smart phone’s dictionary to reduce issues with the spellchecker. When biking and running, speaking audio notes is the easier data entry method. This then creates many audio files that need to be transcribed. While this could potentially be done on device, there is typically not the time while surveying to check and correct on-device transcription results. Instead, we have found that the most efficient workflow is to transcribe the recordings later at a computer.

We use Amazon’s AWS Transcribe speech-to-text web service, since it offers many English accent options, it takes a vocabulary file of common spoken words and phrases, and it allows models to be trained on provided data (we update this regularly with all our manually checked data).

The results from AWS Transcribe are good, although not 100% accurate, especially for noisy field recordings. We maintain a database of common transcription errors and use an R script to automatically clean the AWS Transcribe results, and then we manually check for remaining errors.

### Data processing

We use R scripts with regular expressions to translate the shorthand and spoken observations into spreadsheet format. Use of consistent formatting and consistent species names is essential.

### Case study

Dr. Jon Sullivan has applied this method to a 16.2 km road route between southwestern ōtautahi-Christchurch city and Lincoln town in Canterbury, New Zealand (Fig. 1). This route begins in a suburban landscape of mostly single storey dwellings, with speed limits 50–60 km/hr. It then moves through pastoral farmland, with speed limits 80–100 km/hr, before entering the suburban landscape of Lincoln town with a 50 km/hr speed limit. Over the past two decades, some farmland has been converted into housing and traffic density has consequently increased.

**Figure 1.**
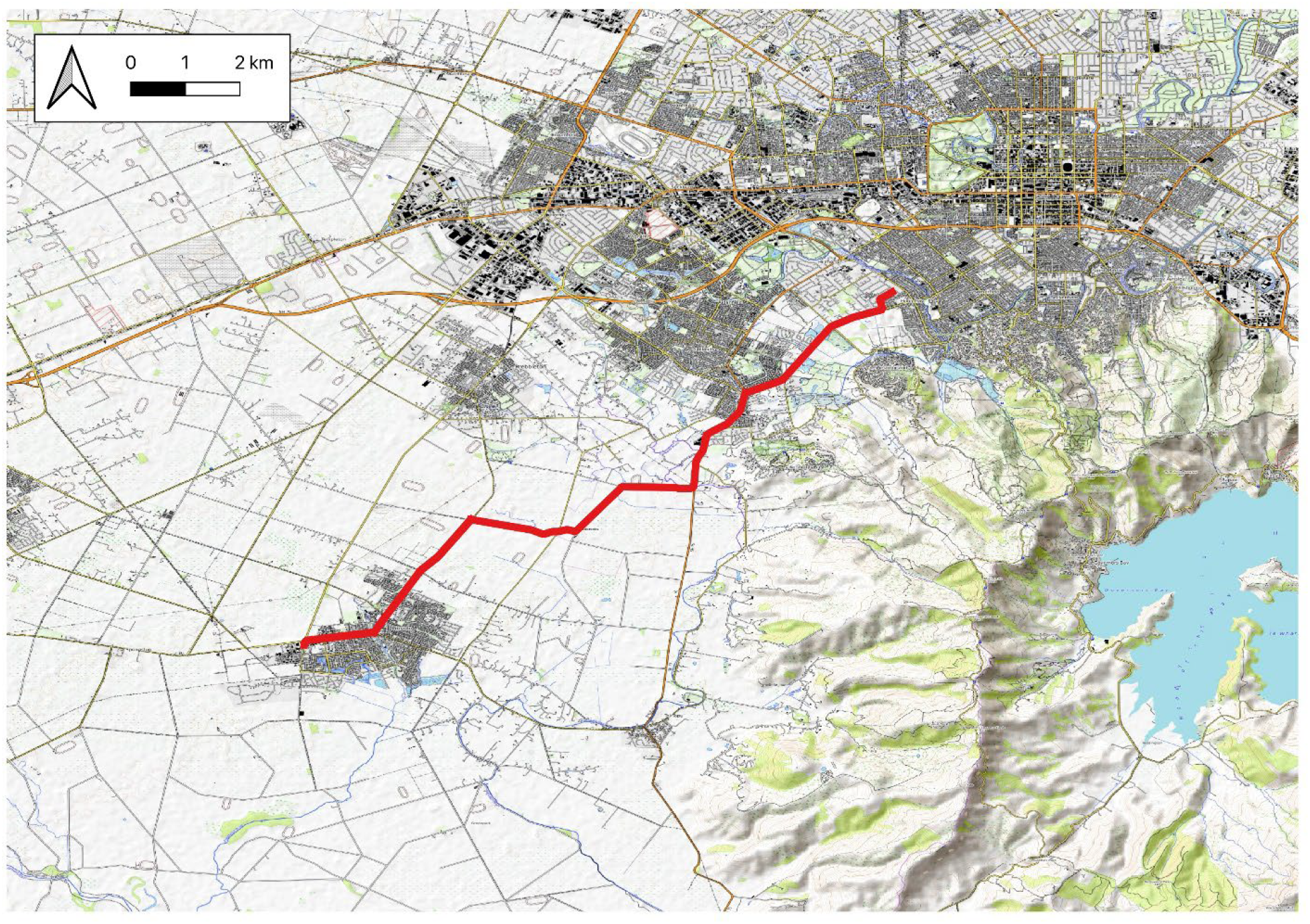
A map of the 16.2 km road route between Christchurch city and Lincoln town where road-kill data were collected from a bike. The area between the suburban housing of Christchurch and Lincoln is primarily pastoral farming. To the southeast are the Port Hills and Banks Peninsula. The background map layer is from Open Topo Map (CC-BY-NC-SA 4.0 OpenTopMap).

Christchurch is a temperate coastal city in the rain shadow of the Southern Alps. Temperature typically ranges from infrequent summer highs over 30ºC to winter night-time lows a few degrees below 0ºC. Average annual rainfall is about 650 mm, spread throughout the year. The built area of the city was originally wetland and evergreen forest on part of the extensive flood plain of the Waimakariri River, which is currently held in place north of the city. Christchurch is now primarily suburban housing and industrial areas surrounded by open pasture and cropping. Many species have been introduced, both on purpose and by accident, almost all since European settlement. Many of the most abundant species in the city are now naturalised introduced species. Many indigenous species are locally, if not nationally, extinct and others are rare.

Along this route our method, or a paper-based precursor of it prior to smart phone apps, has been applied on 2,745 trips since March 2003, primarily travelling by bicycle and occasionally as a passenger in a car. For the Results section below, we report on the 1,652 road-kill trips since 2 April 2012, which were the data collected using a smart phone with the complete method described here. We focus here on reporting summary statistics that illustrate the usefulness of this method; a detailed analysis of these data, along with data from other routes surveyed in this landscape, will be included in a subsequent manuscript.

The data presented include 59 surveys with typed shorthand and 1,593 surveys with transcribed spoken notes. Of the 35,010 spoken road-kill observations (including repeat observations of the same carcasses on consecutive surveys), 3,496 (10%) were manually checked and corrected if necessary (most of these were observations that our R transcription correcting script flagged as needing checking). The remainder are currently as transcribed by AWS Transcribe then automatically corrected with program R using a database of 13,805 transcription errors (these are errors for both road-kill observations and more broadly for other wild counts made along this route). After this automated error correction, there is a 96.7% per character accuracy in the transcriptions, comparing manually checked observations with pre-checked observations.

## Results

### Road-kill taxa

Between 2 April 2012 and 8 March 2024, along one 16.2 km road route, we recorded 4,034 new road-kill carcasses (Table 1). Of the reported road-kill, 62.4% were birds, 31.1% were mammals, and 3% were butterflies. Of the bird road-kill, 18.2% were native species, which reflects the exotic-dominated bird community in this rural and urban landscape. In general, the commonly encountered road-kill were of the most abundant taxa in the surrounding landscape (J. Sullivan, unpublished data).

**Table 1.**
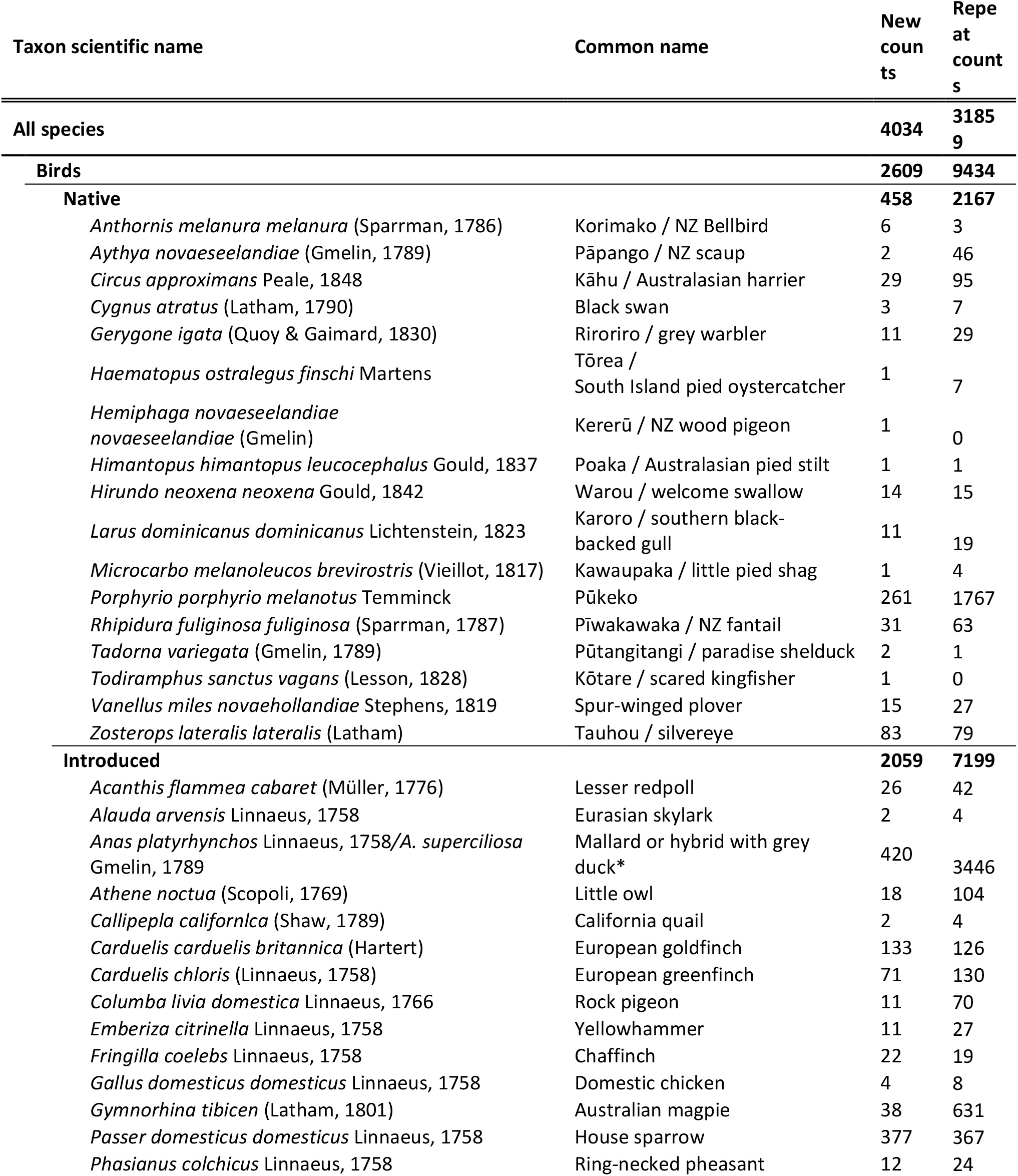
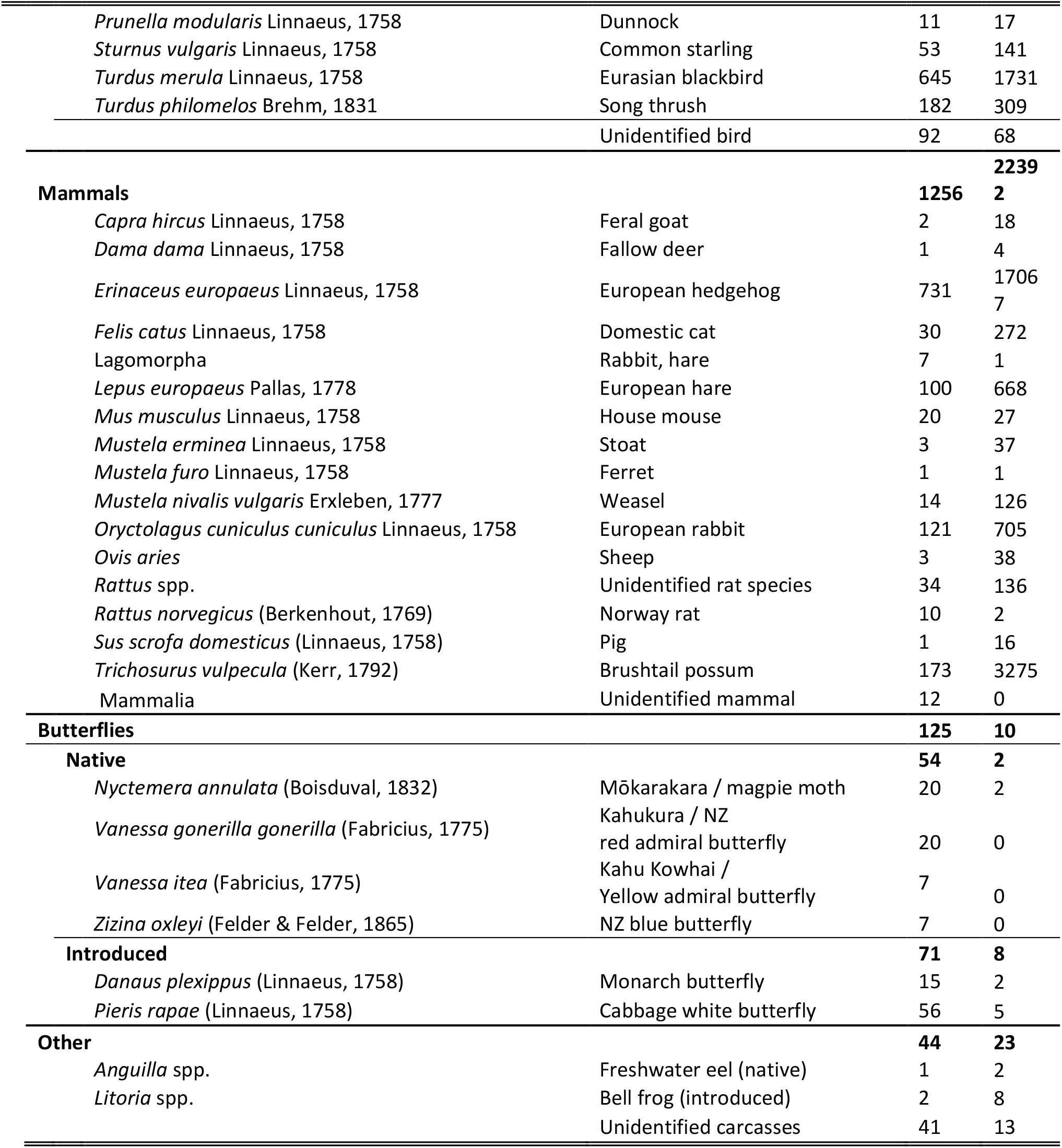
Summary of all road-kill observations recorded in the ten years between 2 April 2012 and 8 March 2024 along one 16.2 km road route. Taxonomic authorities are provided for taxa identified to species level or below. Note that native birds include some recently self-introduced (without human assistance) bird species. All mammalian species in this table are non-native to New Zealand. For the purposes of the survey, the colourful day-flying moth, *Nyctemera annulata*, was consistently included with butterflies. *Note that the mallard and mallardgrey duck hybrids are listed here as introduced species although grey ducks are native.

Figure 2 shows the numbers of the most commonly recorded species. We plot only fresh carcasses, not newly observed carcasses, to avoid possible bias from the taxon variation in carcass persistence. Figure 2 is therefore an underestimate, unbiased by varying carcass persistence, of the number of each species killed during the survey, and so gives a relative measure of how many of each species were killed. The five most frequently killed species on this road route were, in decreasing order, European hedgehog, Eurasian blackbird, mallard/grey duck (mostly hybrids), house sparrow, and pūkeko (see Table 1 for taxon details).

**Figure 1.**
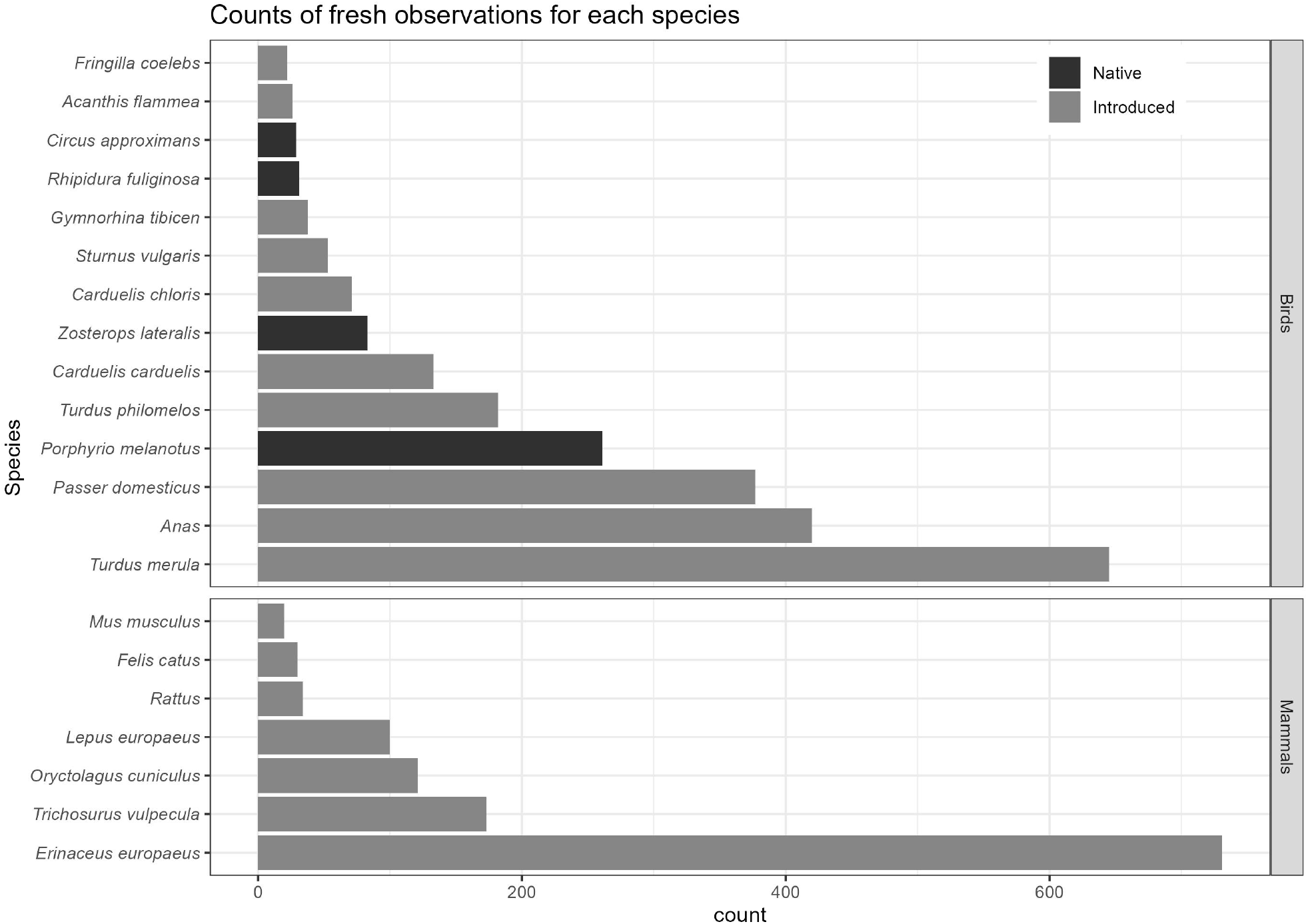
Count of fresh carcass (<24h) observations, for avian and mammalian species with at least 20 fresh carcass observations recorded over the course of 12 years. See Table 1 for the complete taxon names.

**Figure 2.**
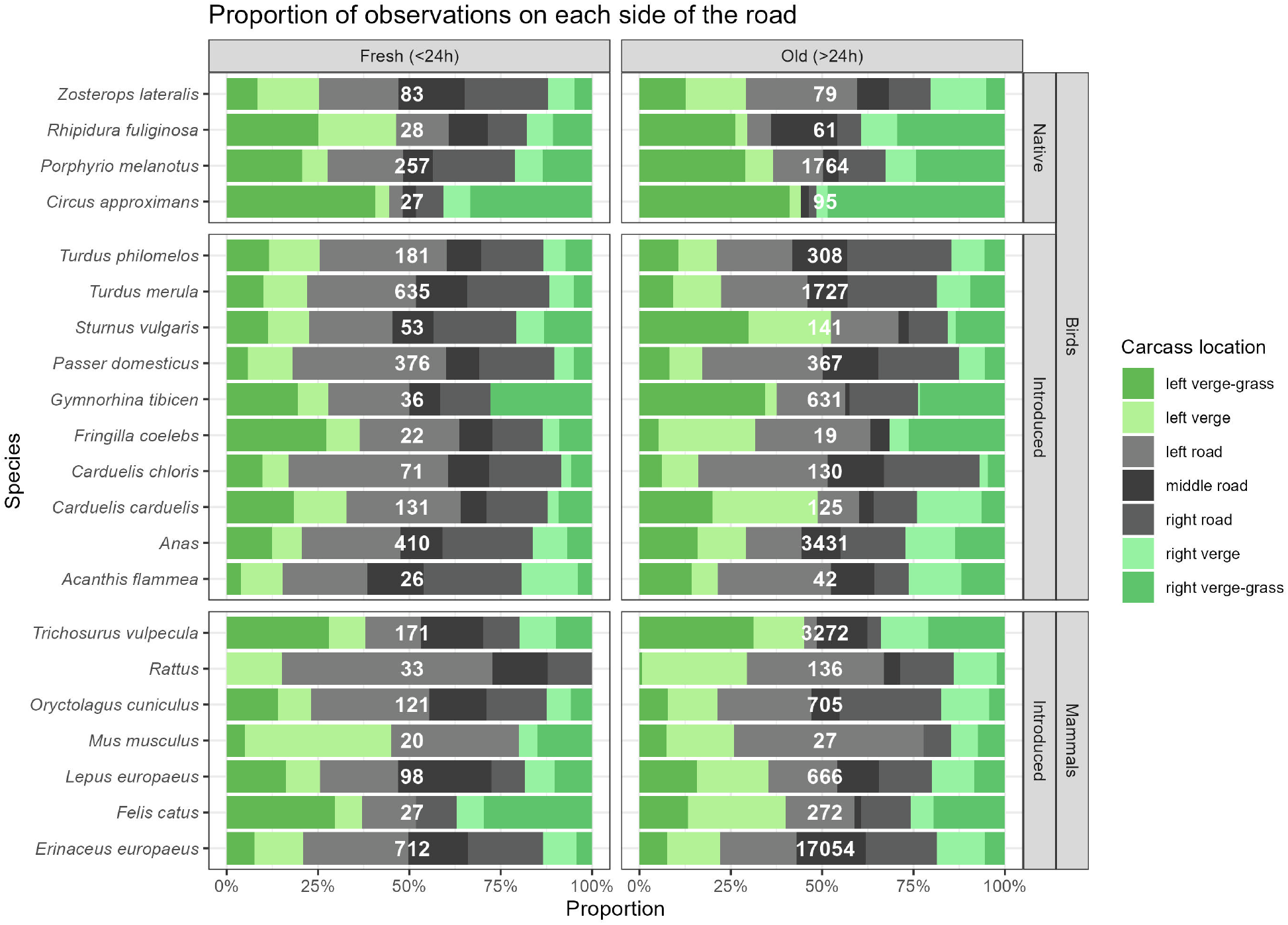
Proportion of carcasses seen on each side of the road, for the avian and mammalian species with at least 20 fresh observations recorded over the course of 12 years. The white labels in the middle of each bar represent the total count of fresh, or old, observations of each species for which a detailed location was recorded.

### Road-kill age

Of the new carcasses, 1,766 (44%) were definitely fresh (<24 hours old) at the time of first sighting, while 778 (19%) were possibly fresh (age “unsure”), and the remainder were definitely >24 hours old. Of all new road-kill, 873 (20%) were definitely >24 hours old and had been missed on the survey the previous day.

### Road-kill condition

Because the route was being surveyed frequently, when road-kill was first reported 17.3% of carcasses were intact and 13.8% had their body cavity exposed but were not yet squashed. Squashed carcasses accounted for 38.3% of newly reported road-kill. As an indication that some new road-kill must have disappeared between surveys and so gone undetected, 22% of road-kill was a fragment of a carcass when first seen, and 1.5% had no body tissue remaining except feathers or fur. Very occasionally black-backed gulls (*Larus dominicanus dominicanus*) and Australasian harriers (*Circus approximans*) were seen removing new carcasses from the road or roadside. Of the repeat reports of old carcasses (“usual” carcasses), 59% were fragments and 21.4% were squashed, with only 3.6% intact and 1.7% exposed.

### Road-kill position

Of new road-kill, 59% was seen on the road (left-road, mid-road, or right-road), 20% was seen on the paved verge of the road, and 21% was on the land beside the road (“grass” in our shorthand). There was considerable variation among taxa in where on and by the road most road-kill was found (Fig. 3). Australasian harriers, in particular, were almost always seen off the road surface (Fig. 3). These are a large-bodied local scavenger that are typically hit as they fly off the road. In contrast, European hedgehogs were mostly seen squashed onto the road surface, as they are slow-moving and defensively roll up when threatened.

**Figure 3.**
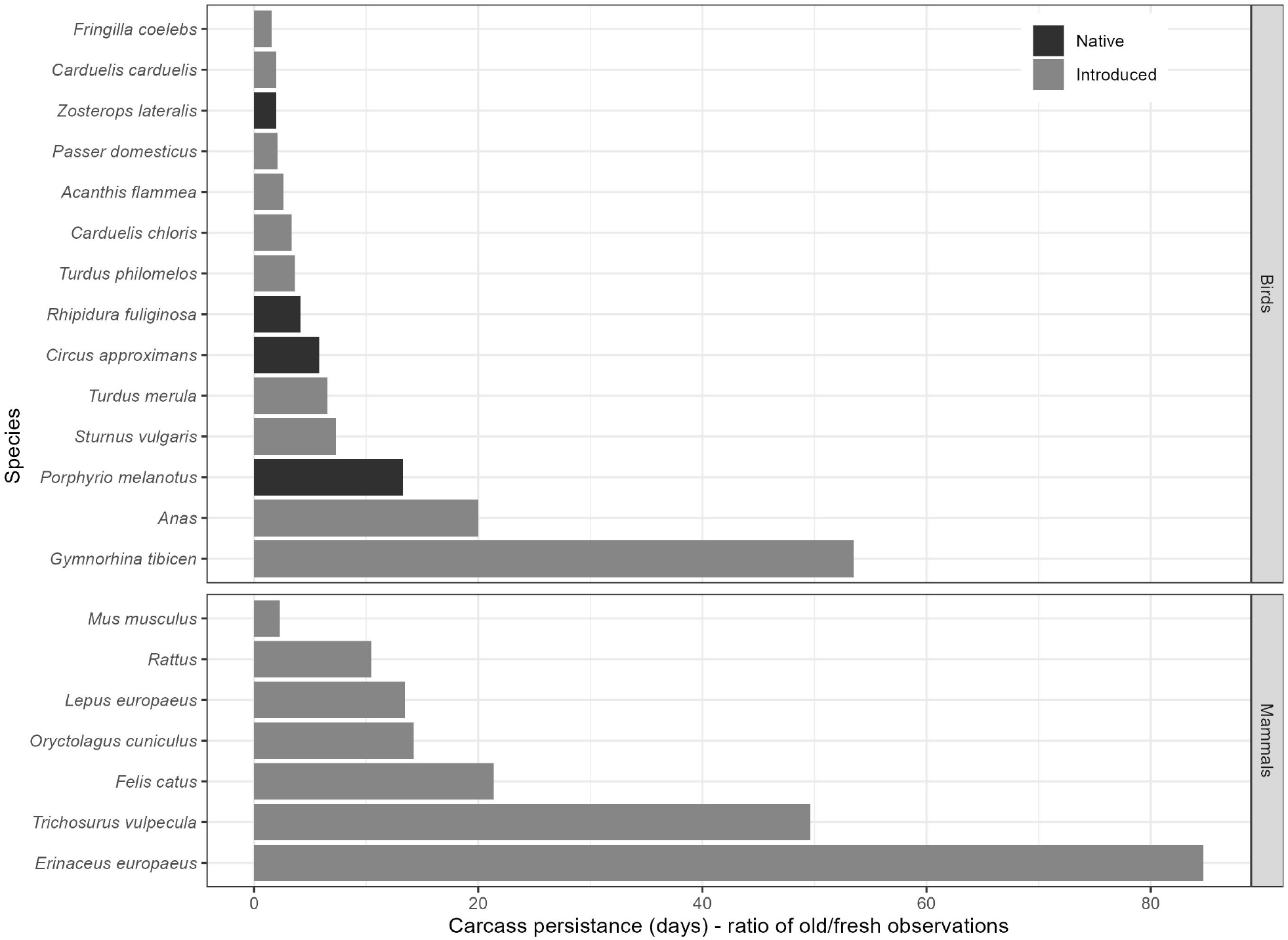
Carcass persistence for avian and mammalian species observed along 16.2 km of road for the avian and mammalian species with at least 20 fresh observations recorded over the course of 12 years. Carcass persistence was calculated as the ratio of fresh (<24h since road-kill) to old (>24h) sightings.

Hedgehogs have tough skin and high persistence and so the ratio of carcasses on and by the road varies little between fresh and old carcasses (Fig. 4). In contrast, large birds like pūkeko (*Pophyrio pophyrio melanotus*) and mallards/grey ducks (*Anas platyrhynchos/A. superciliosa*) had lower persistence on the road and older carcasses were disproportionately found off the road where persistence was longer.

**Figure 4.**
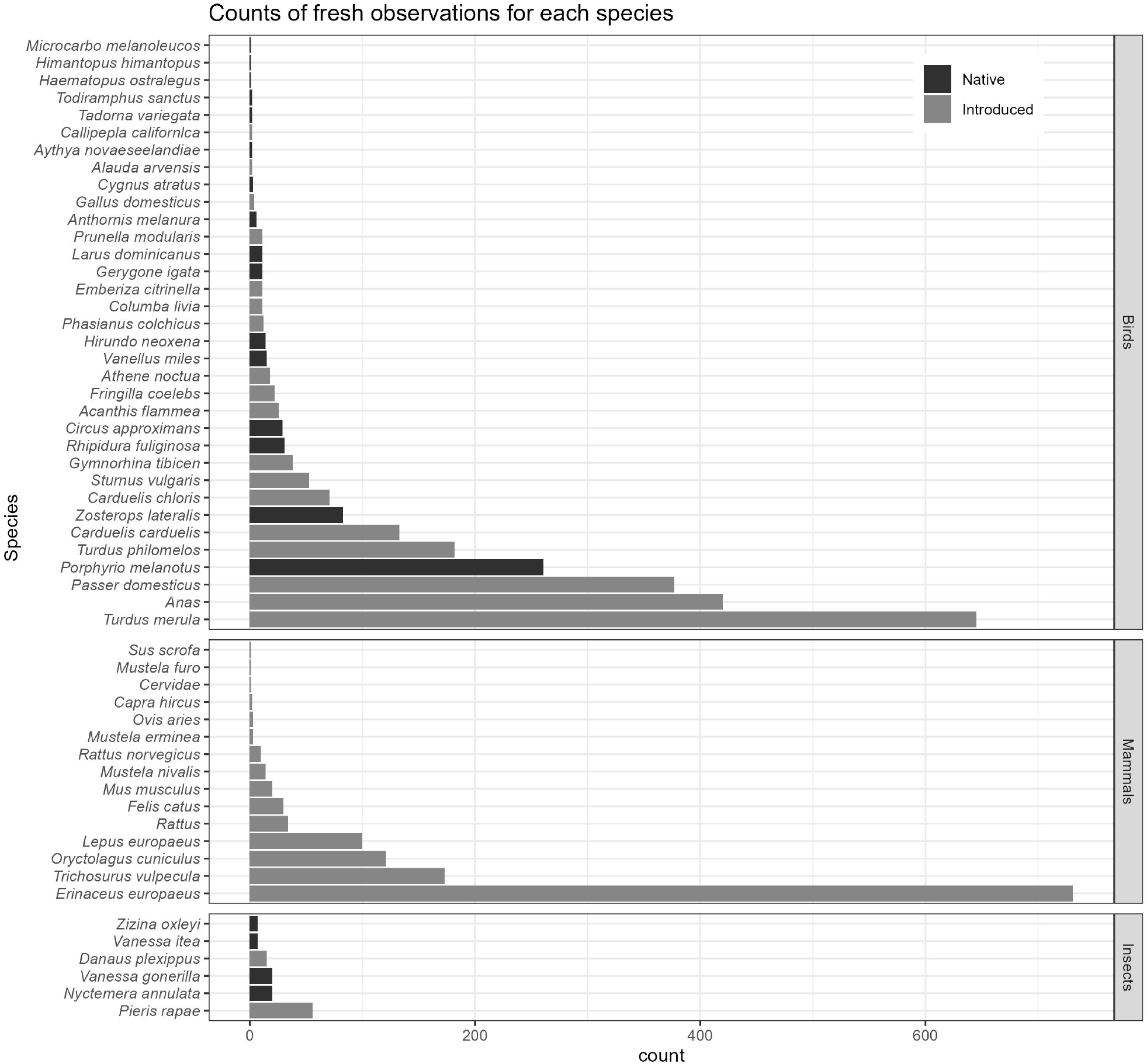
Count of fresh carcass (<24h) observations, for all avian and mammalian species with fresh carcass observations recorded over the course of 12 years.

### Road-kill persistence

There were on average 35.0 road-kill recorded per day (s.d. 23.7, N = 1025 days), with 11.6% of these being new and the rest being repeated observations of older carcasses. Road-kill of some taxa persisted for many months longer than others (Fig. 4). The longest lasting taxon was the European hedgehog, which on occasion lasted on the road for over 12 months. In comparison, small birds typically lasted for only a few days (Fig. 4). This variation in persistence has a big influence on the road-kill present at any one time. While 66.3% of new carcasses were birds, only 33.2% of all carcasses were birds, due to the longer average persistence of mammal road-kill along this route.

### Road-kill rate

These data can be combined to estimate a total road-kill rate for this section of road. On average there were 1.74 new, fresh road-kill carcasses recorded per day over this 16.2 km route (s.d. = 1.82, N = 1652 surveys), and 0.66 new carcasses per day possibly but not certainly <24 hours old (s.d = 1.17). This gives an average estimate of 638.3–880.2 animals killed and detected per year. In addition, since up to 20% of carcasses were missed on the first day, that adds a further up to 180 animals killed per year. When combined, this equates to 3,940–6,544 road-kill/100 km/year. This will still be an underestimate as it necessarily omits animals that were killed then removed by scavengers between surveys, and those that died away from the road surface in out-of-view locations.

## Discussion

There are benefits of the same observers repeatedly surveying the same routes for road-kill, and the method we describe is a practical way of doing this. Repeatedly recording existing carcasses provides information on the persistence of carcasses. We found that carcass persistence can vary considerably among taxa, with tough-skinned taxa like hedgehogs often remaining on the road for many months (and sometimes years) while smaller and weaker skinned taxa would be gone in days to weeks. This needs to be taken into account when interpreting less frequently recorded road-kill data, as taxa with more persistent carcasses will be over-represented in single road-kill surveys. Repeat surveys can also reveal the seasonality of road-kill, and effects of varying weather and traffic volumes on road-kill and its persistence (Erritzoe, Mazgajski & Rejt, 2003; Guinard, Julliard & Barbraud, 2012; van Der Ree, Smith & Grilo, 2015).

A more complete picture of road-kill is also gained by consistently including the location of each carcass on and by the road. In our example, different taxa were typically found on different parts of the road and road verge (e.g. Australasian harriers were almost always on the grass verge while hedgehogs were almost always on the road surface). Detection probability will generally be higher on the road surface, where carcasses are more visible, than by the sides of the road (something we plan on modelling with our data in a future manuscript). Road-kill persistence also differs based on road location. Both of these issues mean that road-kill data can contain biases in the taxa detected when road-kill location is not accounted for.

An example of these issues can be found in Brockie et al. (2009), New Zealand’s largest published road-kill study to date. The length of New Zealand’s North Island was surveyed for road-kill from a car once a decade for three decades. The majority of the road-kill carcasses in each decade were hedgehogs and brush-tailed possums, with very few small birds seen dead. Our South Island survey data suggests that this result is due to the long persistence of hedgehog and possum carcasses on the road relative to smaller birds (Fig. 4), combined with car-based surveys missing a lot of smaller birds dead in the roadside grass.

Data from frequent, standardised repeat surveys of road-kill using our method, or similar, will allow for per-taxon carcass persistence and detection probability to be estimated. By using the method with a mix of transport modes along the same route (e.g. walking, running, biking, passenger in a car), it should also be possible to calculate detection probabilities for different transport modes. This is analogous to distance sampling methods (Thomas et al., 2010), where it is assumed that animals at distance zero are always detected and statistics can estimate the decline in detection with distance. The equivalent should be possible for road-kill, assuming that carcasses are always detected while walking and detection probability will decline with observer speed (doing this will require a more in-depth analysis of our data than we present here). When persistence and detection probability have been well estimated for an area, it will be possible to calibrate the results of more infrequent surveys, like the per decade survey of Brockie et al. (2009). For example, if the ratio of birds to mammals in our South Island survey (62%) is comparable to the North Island, we estimate that Brockie et al. (2009) detected <9% of the bird carcasses in their survey.

Most literature on road-kill reports on studies that tallied road-kill per unit distance, rather than geotagging individual carcasses. For example, Brockie et al. (2009) recorded road-kill locations using the vehicle’s odometer to the nearest kilometre. Valuable data is lost in this simplification, such as identifying areas of the road, and habitats, that are hotspots for road-kill. Our method demonstrates that this simplification is no longer necessary. In this age of smart phones, it is just as simple for the observer to geotag each carcass when observed. Such data can always be tallied up per unit distance later if needed to compare with past survey results.

Our method separates the concepts of whether a carcass is a new or repeat observation, from the carcass age (<24 hours old, uncertain, or >24 hours old). We see value in both. Consistently recording whether a carcass is new, or a repeat of a previously recorded carcass (“usual” carcasses), is essential for documenting carcass persistence. In our case, where one observer is traveling the same route almost daily, memory and a GPS are adequate for tracking which carcasses are new and which are usual. When survey frequency is reduced, and/or multiple observers are used, it may be beneficial to stop and mark carcasses directly. However, doing so will add time and complexity to a survey and we anticipate that in most cases a taxon identification and GPS coordinate should be sufficient to know whether a carcass is new or usual. This repeated recording of the same carcasses also has potential for estimating detection probability, by assessing how many repeat carcasses are missed in surveys.

Estimating carcass age is helping us to assess detection probability (such as when a clearly old carcass is detected for the first time on a route that was recently surveyed, indicating the carcass was previously missed). When combined with carcass persistence, we also expect this will help us to estimate how many carcasses may be killed then removed by scavengers before we record them. For less frequently surveyed routes, this ratio of fresh to old carcasses can be valuable for calibrating biases due to taxon-variable carcass persistence.

The overall road-kill rate in our case study is estimated at 3,940–6,544 road-kill/100 km/year. This is near the high end of international estimates (Calvert et al., 2013) and higher than previous results from previously published New Zealand road-kill studies (Brockie, Sadleir & Linklater, 2009; Sadleir & Linklater, 2016; Flux, Tryjanowski & Zduniak, 2022). The estimates collated by Calvert et al. (2013) ranged from 487 to 37,280 road-kill/100 km/year with a median of 1,488 road-kill/100 km/year. Our estimate, while higher than this median, will be an underestimate as some carcasses will have been missed or removed by scavengers before they could be detected. The combination of surveying by bike plus repeated surveys of carcasses likely make our estimate closer to the true rate than from car-based and less frequently travelled surveys. Collectively, these studies emphasise the role of road-kill as a potentially important source of mortality for species occurring in landscapes characterised by dense road infrastructure and high vehicle speeds.

## Conclusions

The monitoring method described here provides a useful step toward standardised data collection on wildlife mortality along roads. Its versatility and adaptability make it suitable across a wide range of landscapes, habitats, road types, and species, offering broad utility for researchers and conservation practitioners. Relying on readily available technologies such as GPS enabled smartphones and speech to text functionalities, the method allows efficient and accurate recording of carcass locations and associated metadata without requiring observers to stop. Specifically, it captures precise location data through continuous GPS tracking and detailed metadata through speech to text, enabling rapid data collection during regular travel by walking, cycling, or as a passenger in a vehicle.

While our case study focused on a specific landscape and survey context, the method can be readily adapted for other scenarios. It may be applied to different regions, habitats, or target species, and tailored to address specific research or conservation questions. For example, if temporal variation is a key factor but data on carcass persistence are already available, the method could be modified to sample less frequently and instead focus on covering greater spatial extent or finer seasonal resolution. This approach may be particularly effective for larger species, such as kiwi or penguins, which are likely more easily detected and persist longer on the road surface than passerines. The method also allows for incorporation of updated procedures or new technologies as they become available, ensuring continued relevance in future applications.

Data generated through this approach can support analyses at multiple ecological scales and help reveal patterns in carcass persistence and detection probability, both of which are essential for understanding road-kill impacts. The comprehensive collection of georeferenced data enables identification of road-kill hotspots and provides valuable baseline information for assessing the effectiveness of mitigation efforts. Although the method is manual, it uses accessible technologies in ways that reduce cost and increase flexibility, making it efficient and easy to incorporate into daily routines.

By establishing consistent data collection parameters, this method addresses several important sources of bias that have limited previous road-kill research. This improves the comparability and reliability of data across studies. The approach outlined here offers a practical and scalable model for systematic monitoring of wildlife mortality on roads, supporting targeted conservation measures, assessments of population level impacts, and informed mitigation planning within New Zealand and in other regions worldwide.

## Acknowledgements

We thank John Innes for his valuable insight on New Zealand avifauna and the New Zealand Transport Agency Waka Kotahi for feedback on earlier discussions on impacts of road-kill.

## References

Beauchamp A. 2009. Bird deaths on riverside drive between Whangarei and Onerahi, New Zealand. Notornis 56:95–97.

Brockie R. 2007. Notes on New Zealand mammals 4. Animal road-kill “blackspots.” New Zealand Journal of Zoology 34:311–316. DOI: 10.1080/03014220709510090.

Brockie RE, Sadleir RMFS, Linklater WL. 2009. Long-term wildlife road-kill counts in New Zealand. New Zealand Journal of Zoology 36:123–134. DOI: 10.1080/03014220909510147.

Burger J, Gochfeld M. 1992. Vulnerability and mortality of young Australian Magpies on roads. The Wilson Bulletin 104:365–367.

Calvert AM, Bishop CA, Elliot RD, Krebs EA, Kydd TM, Machtans CS, Robertson GJ. 2013. A Synthesis of Human-related Avian Mortality in Canada. Avian Conservation and Ecology 8:art11. DOI: 10.5751/ACE-00581-080211.

van Der Ree R, Smith DJ, Grilo C (eds.). 2015. Handbook of Road Ecology. West Sussex, UK: John Wiley & Sons Ltd. DOI: 10.1002/9781118568170.

Erritzoe J, Mazgajski TD, Rejt Ł. 2003. Bird casualties on European roads — a review. Acta Ornithologica 38:77–93. DOI: 10.3161/068.038.0204.

Flux JEC. 2019. Swamp harrier (Circus approximans) road-kills, 1962-2018, and the effect of rabbit density. Notornis 66:217–220.

Flux JEC, Tryjanowski P, Zduniak P. 2022. Road-kills in New Zealand: long-term effects track population changes and reveal colour blindness. European Journal of Ecology 8:30–42. DOI: 10.17161/eurojecol.v8i2.18567.

Freeman S. 2010. Western weka road-kill at Cape Foulwind, Buller, New Zealand. New Zealand Journal of Zoology 37:131–146. DOI: 10.1080/03014223.2010.482972.

Guinard É, Julliard R, Barbraud C. 2012. Motorways and bird traffic casualties: Carcasses surveys and scavenging bias. Biological Conservation 147:40–51. DOI: 10.1016/j.biocon.2012.01.019.

Guinard É, Prodon R, Barbraud C. 2015. Case study: a robust method to obtain defendable data on wildlife mortality. In: Van der Ree R, Smith DJGC ed. Handbook of Road Ecology. West Sussex, UK: John Wiley & Sons Ltd, 96–100. DOI: 10.1002/9781118568170.ch12.

Huijser MP, McGowen P, Fuller J, Hardy A, Kociolek A, Clevenger AP, Smith D, Ament R. 2008. Wildlife-vehicle collision reduction study. Report to Congress. Wasington DC, USA: U.S. Department of Transportation, Federal Highway Administration.

Husby M. 2017. Traffic influence on roadside bird abundance and behaviour. Acta Ornithologica 52:93–103. DOI: 10.3161/00016454AO2017.52.1.009.

Pagany R. 2020. Wildlife-vehicle collisions - Influencing factors, data collection and research methods. Biological Conservation 251:108758. DOI: 10.1016/j.biocon.2020.108758.

Robertson HA, Baird KA, Elliott GP, Hitchmough RA, McArthur NJ, Makan TD, Miskelly CM, O’Donnell CJF, Sagar PM, Scofield RP, Taylor GA, Michel P. 2021. Conservation status of birds in Aotearoa New Zealand, 2021. Wellington, New Zealand.

Rytwinski T, Soanes K, Jaeger JAG, Fahrig L, Findlay CS, Houlahan J, van Ree RD, van Der Grift EA. 2016. How effective is road mitigation at reducing road-kill? A meta-analysis. PLoS ONE 11:1–25. DOI: 10.1371/journal.pone.0166941.

Sadleir RMFS, Linklater WL. 2016. Annual and seasonal patterns in wildlife road-kill and their relationship with traffic density. New Zealand Journal of Zoology 43:275–291. DOI: 10.1080/03014223.2016.1155465.

Thomas L, Buckland ST, Rexstad EA, Laake JL, Strindberg S, Hedley SL, Bishop JRB, Marques TA, Burnham KP. 2010. Distance software: design and analysis of distance sampling surveys for estimating population size. Journal of Applied Ecology 47:5–14. DOI: 10.1111/j.1365-2664.2009.01737.x.

Washington C, Paterson A, Sixtus C, Ross J. 2008. Roadside behaviour of Porphyrio porphyrio melanotus (Aves: Rallidae). New Zealand Natural Sciences 33:33–41.

